# Randomization based evaluation of distinct topological and cancer expression characteristics of mutually acting gene pairs

**DOI:** 10.1101/2024.10.09.617344

**Authors:** Ertuğrul Dalgıç, Muazzez Çelebi-Çınar, Merve Vural-Özdeniz, Özlen Konu

**Author notes:** Equal corresponding authors: ED and OK. Equal contribution (Muazzez Çelebi-Çınar, Merve Vural-Özdeniz).

## Abstract

Small scale molecular network patterns and motifs are crucial for systems level understanding of cellular information transduction. A subgroup of such patterns consists of only 2 molecules, which we named as mutually acting pairs. Using randomizations, we statistically explored such gene pair designs, i.e. mutually positive (PP) or negative (NN) and positive-negative (PN) pairs, in two comprehensive and distinct large-scale molecular networks from literature; the human protein signaling network (PSN) and the human gene regulatory network (GRN). While the numbers of NN and PN pairs were significantly higher, the number of PP pairs was significantly lower than randomly expected values. Genes participating in mutual pairs were more connected than other genes. The connectivity of mutual gene pairs differed between the two networks. NN genes were more connected than PP and PN in GRN for all types of degree values, including in, out, positive or negative connections, but less connected for in-degree and more connected for out-degree values in PSN. They also had significantly high number of intersections with each other and PN pairs than randomly expected values, indicating potential cooperative mechanisms. The three mutual interaction designs we examined had distinct RNA and protein expression correlation characteristics. NN protein pairs were uniquely over-represented across normal tissue samples, whose negative correlations were lost across cancer tissue samples. PP and PN pairs showed non-random positive RNA or protein expression correlation across normal or cancer tissue samples. Moreover, we developed an online tool, i.e., MGPNet, for further user specific analysis of mutual gene pairs. For example, we identified SNCA with significantly enriched negatively correlated NN pairs with ubiquitin-proteasome members only in normal tissues. Unique non-random characteristics of mutual gene/protein pairs identified in two different comprehensive human molecular networks could provide valuable information for a better understanding of molecular design principles between normal and cancer states.

## Introduction

Small scale subnetwork patterns have previously been investigated in large scale biological networks such as gene regulatory and protein signaling networks (Shen-Orr et al. 2002, Milo et al. 2002, Yeger-Lotem and Margalit 2003, Mazurie et al. 2005, Alon et al. 2007, Borotkanics and Lehmann 2015). Based on these studies some patterns emerged as being recurrent. Network motifs were defined as the small network patterns that take place significantly in biological networks more than random (Shen-Orr et al. 2002, Milo et al. 2002). Such motifs could be responsible for the information processing in the networks, such as persistence detection, pulse generation, speeding up or slowing down responses (Alon U. 2007). They may also have a significant role in human diseases; specialized motifs could play significant roles for the switch like behavior of biological systems, for instance transition from normal to cancer (Shiraishi et al. 2010, Siegal-Gaskins et al. 2011).

There is a moderate correlation between mRNA and protein expression levels of human protein-coding genes while the significance of correlation varies among genes (Kosti et al. 2016, Prabahar et al. 2024). Protein level changes, which are not reflected at the genomic level, could play important roles in carcinogenesis (Jarnuczak et al. 2021). Therefore, integrative analysis with protein datasets is also necessary for deciphering the systems biology of cancer. Two major types of molecular cellular networks, i.e., gene regulatory (transcriptional) networks (GSN) and protein signaling networks (PSN), were both previously analyzed for motifs but not extensively compared with each other in human (Shen-Orr SS et al. 2002, Alon U. 2007, Shiraishi et al. 2010, Siegal-Gaskins et al. 2011, Borotkanics and Lehmann 2015).

Motif analyses in gene or protein level networks have dominantly focused on patterns with at least 3 members (Shen-Orr SS et al. 2002, Alon U. 2007, Defoort et al. 2018, Zenere et al. 2021). A few studies statistically analyzed 2-node (pairwise) structures, but they omitted the sign of interactions (activation or inhibition) and did not investigate simultaneously the two different types of human molecular network datasets (Mazurie et al. 2005, Wang et al. 2014, Borotkanics and Lehmann 2015). Unlike bigger and more complex patterns of network structures, simple mutually acting gene pairs (aka. mutual gene pairs) were not extensively analyzed in large scale signed human molecular networks. Here we focused on these mutually positive (PP), mutually negative (NN) and positive-negative (PN) pairs and provided evidence for their distinct characteristics that might hint to specific roles for information transmission in the cellular molecular networks. In particular, NN pairs showed distinct connectivity and expression correlation characteristics. Any gene or genes of interest could be specially analyzed for their involvement in mutual pair networks by the Shiny R based online tool MGPNet we developed.

## Results

### Mutual Gene Pairs

There are only three possible pairwise patterns for mutual gene pairs in signed directed networks; mutually positive (PP), mutually negative (NN) and positive-negative (PN) pairs (Figure 1A). We analyzed the number of mutual gene pair patterns in two separately comprehensive human molecular networks, namely, the gene regulatory network (GRN) and the protein signaling network (PSN). GRN was constructed by the integration of various different resources (Müller-Dott S et al. 2023). PSN was built by the integration of curated human cancer signaling networks with the BIOGRID protein-protein interactions (Awan et al. 2007; Cui et al. 2007; Li et al. 2012, Wang et al. 2014). GRN is based on transcriptional interactions for no particular type of sample whereas PSN is based on signaling pathways of cancer. We investigated 2-node signed subnetwork designs in GRN and PSN, because they are the two most comprehensive large scale public human molecular networks, yet they share less than 10% of their connections. Thus, they are two distinct signed directed molecular networks.

**Figure 1.**
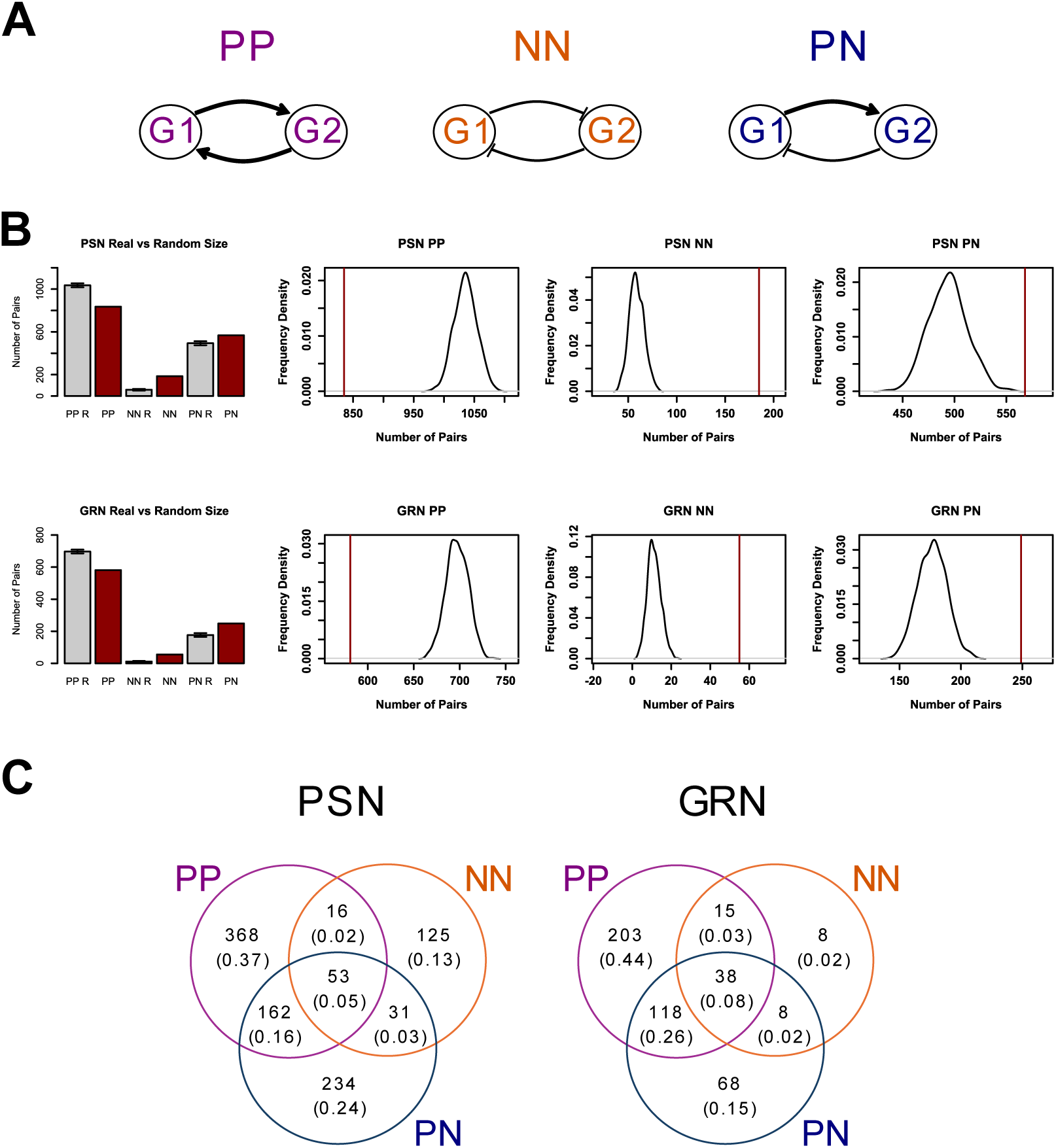
Types of mutual gene pairs and their distributions in large scale human molecular networks. A. Three different mutual pair patterns: mutual positive (PP), mutual negative (NN) and positive-negative (PN). B. Number of PP, NN, and PN pairs (red) in PSN and GRN with comparison to the kernel density plots of randomly expected numbers (R; gray). C. Overlap between the genes belonging to different types of mutual pairs in both networks was shown as a Venn diagram. Besides the actual number of genes, the fraction of the genes over the union of all genes participating in any mutual pairs was shown in parentheses.

In both GRN and PSN, the number of NN or PN pairs we extracted were significantly higher, whereas the number of PP pairs were significantly lower than random (random sampling p-value < 0.001) (Figure 1B). Therefore, NN and PN patterns are network motifs in two different types of human molecular networks whereas PP pattern is not. The number of genes participating in NN pairs were lower than PP or PN pair genes in both networks (Figure 1C). The overlap between the PP and PN pair genes were higher than their overlap with NN pairs. In contrast to PP and PN pair genes, NN pair genes did not include a high fraction of unique genes that did not participate in other mutual pairs (Figure 1C, 0.13 vs 0.24-0.37 in PSN and 0.02 vs 0.15-0.44 in GRN).

### Connectivity of Mutual Gene Pairs

To evaluate the connectivity characteristics of mutual gene pairs different types of degree values were analyzed depending on the type of interactions. In both the entire signed directed networks, total degree values of the genes that participated in mutual pairs were higher than the remaining genes with at least two connections (Figure 2, two-sided Wilcoxon test p-value < 2.2e-16). NN pair genes were more connected than PP and PN pair genes in GRN, but not in PSN (one-sided Wilcoxon test p-value < 0.0005 for GRN, 0.87 for PSN).

**Figure 2.**
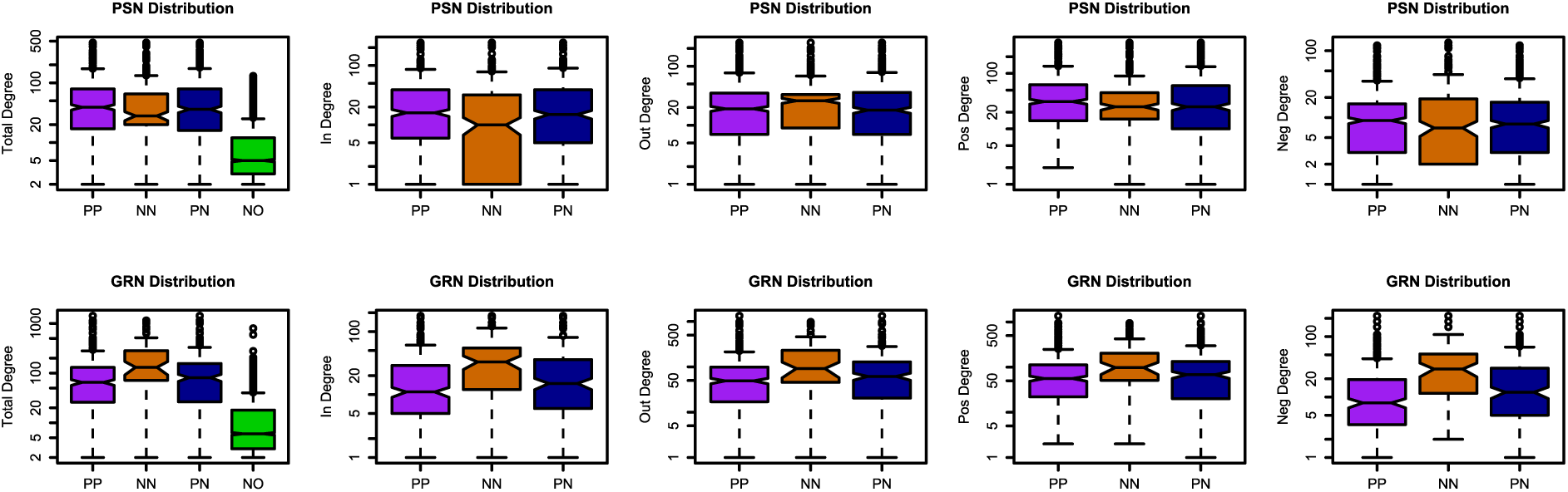
Degree values of mutual pair genes (PP, NN, and PN) and nonmutual pair genes (NO) in PSN and GRN were shown by boxplots. Distribution of total, in, out, positive and negative degree values were compared for the mutual pair genes. Nonzero values were plotted in log scale. NO genes were limited to those with at least two interactions.

Next, we considered other types of degree values, i.e., in/out and positive/negative. In-degree values of NN pairs were lower than PP and PN pairs while out-degree values were higher in PSN (Figure 2, one-sided Wilcoxon test p-value < 0.0001 for in-degree, 0.03 for out-degree). No consistently significant difference was observed for positive- or negative-degree. The cause for the lower in-degree values of NN pair genes was mostly based on the finding that almost half (58 of 125) of uniquely NN pair genes had only one in-degree value (Supplementary Table 1). On the other hand, SNCA, MYB, UBE3A, and CDK6 were the top ranking uniquely NN pair genes with high out-degree values (Supplementary Table 1).

Unlike PSN, NN pairs were more connected than PP and PN for all types of degree values in GRN (Figure 2, one-sided Wilcoxon test p-value < 0.003). Overall, NN pair genes had different degree values in comparison to other mutual pairs in PSN and GRN. Since the genes belonging uniquely to NN motif was low in GRN (8 out of 69 NN pairs) and they did not have high degree values, and hence the higher connectivity of NN was mostly due to the high degree values of the genes that overlap with PP and/or PN patterns (Figure 1C and 2, Supplementary Table 1). Some of these highly connected genes were TP53, MYC, RELA, NFKB1, which had all 3 types of mutual relationships (Supplementary Table 1).

Next, we analyzed the intersection of mutual pairs, i.e., for how many occurrences two different mutual pairs share a common gene (Figure 3A). In both GRN and PSN, the number of NN-NN or PN-PN co-occurences were significantly higher, whereas the number of PP-PP co-occurences were significantly lower than random (random sampling p-value < 0.001) (Figure 3B). In addition, the number of PN-NN intersections were also significantly higher in both networks (random sampling p-value < 0.001), whereas no common significant trend was observed for PP-NN or PP-PN intersections (Figure 3B).

**Figure 3.**
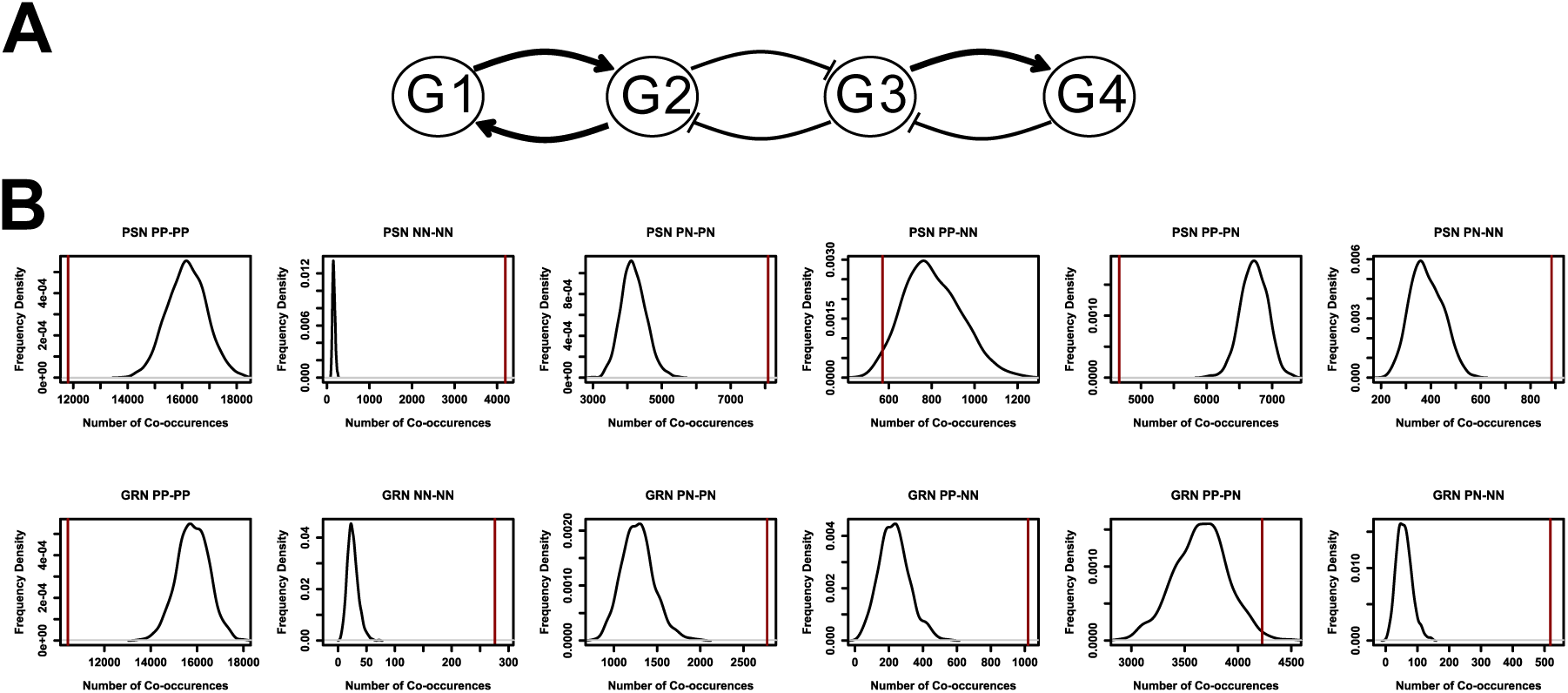
Co-occurences of mutual pair patterns in PSN and GRN. A. Co-occurrences of PP, NN and PN motifs were shown in a sample network, where there is a single co-occurrence of PP vs NN patterns via G2, a single co-occurrence of PN vs NN motifs via G3. B. Number of co-occurences for PP vs PP, NN vs NN, PN vs PN, PP vs NN, PP vs PN, and PN vs NN (red line) were compared to the kernel density plots of randomized co-occurence values (black).

### RNA Coexpression in Normal vs Cancer Tissues

For evaluation of the role of mutual gene pairs in cancer, our approach was to integrate RNA or protein level expression values obtained from Human Protein Atlas (HPA), specifically for normal or cancer tissue samples, so that coexpression between the mutual pair genes could be analyzed comparatively (see Methods). We explored the RNA expression correlation of PP, NN and PN pairs for both GRN and PSN in cancer vs normal tissue samples together with non-mutual pairs (NO).

NN pairs did not have a significant difference of coexpression values in normal tissues for PSN when compared to PP, PN and NO pairs, or to the remaining gene pairs with a negative interaction (gene pairs with a negative interaction but no participation in any NN motif) (Figure 4, Kolmogorov–Smirnov test p-value < 0.15, two-sided Wilcox test p-value < 0.582). Further, we compared the median RNA coexpression value of NN pairs to random negatively interacting pair lists of the same size and found no significant difference for lower values (randomization p-value = 0.662). In contrast, NN pairs had significantly lower coexpression in cancer for PSN (Figure 4, Kolmogorov–Smirnov test p-value < 0.031, two-sided Wilcox test p value 0.018, randomization p-value = 0.031). There was no significant correlation difference of NN pairs for GRN (Kolmogorov–Smirnov test p-value < 0.658, two-sided Wilcox test p-value < 0.593 for normal, Kolmogorov–Smirnov test p-value < 0.741, two-sided Wilcox test p-value < 0.581 for cancer, randomization p values for lower median NN coexpression are 0.826 for normal and 0.68 for cancer). Cancer vs normal tissue differential RNA correlation values of NN pairs were significantly lower than PP, PN or NO pairs in PSN (Kolmogorov–Smirnov test p-value < 0.081, one-sided Wilcox test p-value < 0.033), but not in GRN (Kolmogorov–Smirnov test p-value < 0.831, two-sided Wilcox test p-value < 0.907).

**Figure 4.**
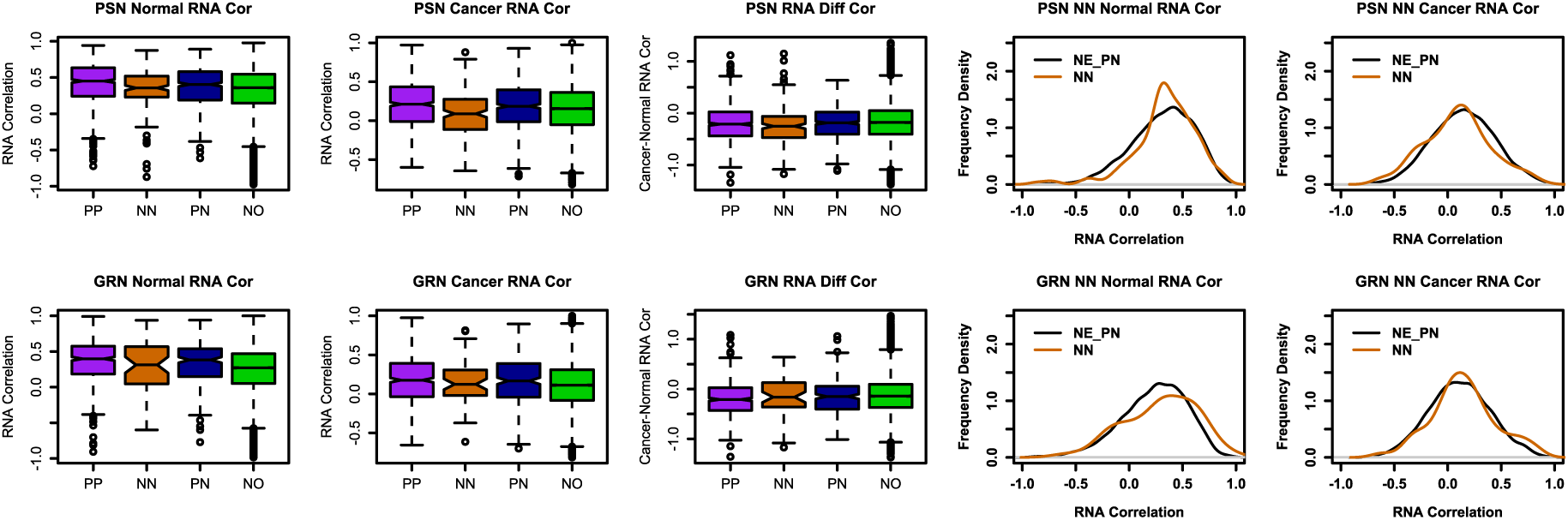
RNA expression correlation distributions of mutual pair genes (PP, NN, and PN) and and nonmutual pair genes (NO) in PSN and GRN were shown by boxplots. Kernel density plots of NN pair correlation values together with the remaining genes with negative interactions; non-mutual negative interaction pairs (NE) and PN pairs. Correlation (cor) was measured by Spearman’s rho. Correlation analysis included normal and cancer tissue specific expression values.

Unlike NN pairs, PP and PN pairs had significantly higher RNA correlation values, when compared with other pairs with positive and/or negative interactions, for both tissues, normal and cancer, in both PSN and GRN (Supplementary Text and Supplementary Figure 1). Thus, NN pairs were unique in their differential RNA coexpression in cancer vs normal tissues, such that they had lower coexpression values in cancer for PSN.

### Protein Coexpression in Normal vs Cancer Tissues

Besides RNA coexpression, we investigated protein level correlation of mutual gene pairs in the same way. NN pairs had significantly lower coexpression in normal but not in cancer for PSN when compared to the PP, PN and NO pairs or the remaining pairs with a negative interaction (Figure 5, Kolmogorov–Smirnov test p-value < 0.002 for normal and 0.847 for cancer, one-sided Wilcox test p-value < 0.0005 for normal and 0.743 for cancer). When compared to random negatively interacting pair lists of the same size, the median protein coexpression value of NN pairs was significantly lower only in normal tissues for PSN (randomization p-value = 0.002 for normal, 0.55 for cancer). On the other hand, they had no significant protein coexpression difference in normal tissues but higher coexpression values than the remaining pairs with a negative interaction in cancer for GRN (Kolmogorov–Smirnov test p-value < 0.716 for normal and 0.155 for cancer, one-sided Wilcox test p-value < 0.352 for normal and 0.02 for cancer, randomization p-value = 0.391 for normal, 0.047 for cancer). Furthermore, NN pairs had greater cancer vs normal differential coexpression values than PP, PN or NO pairs in PSN (Kolmogorov–Smirnov test p-value < 0.037, one-sided Wilcox test p-value < 0.003), but not in GRN (Kolmogorov–Smirnov test p-value < 0.822, two-sided Wilcox test p-value < 0.277).

**Figure 5.**
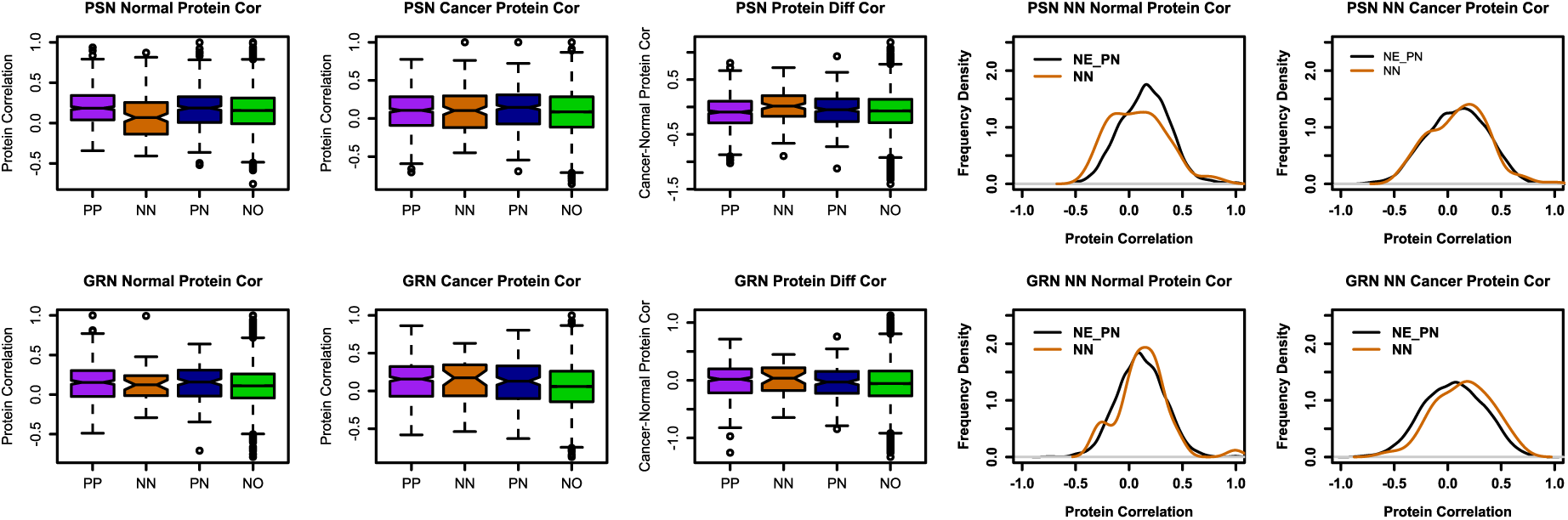
Protein expression correlation distributions of mutual pair genes (PP, NN, and PN) and and nonmutual pair genes (NO) in PSN and GRN were shown by boxplots. Kernel density plots of NN pair correlation values together with the remaining genes with negative interactions; non-mutual negative interaction pairs (NE) and PN pairs. Correlation (cor) was measured by Spearman’s rho. Correlation analysis included normal and cancer specific expression values.

Dissimilar to NN pairs, PP and PN pairs had significantly higher protein correlation values in normal or cancer tissues (Supplementary Text and Supplementary Figure 1). Hence, NN pairs were distinctive in their differential protein coexpression in cancer tissues vs normal, such that they had lower coexpression values in normal for PSN.

### Visualization of Mutual Pairs by the Online Tool MGPNet

We have constructed an online network analysis tool, which we named MGPNet, in order to aid the visualization and exploration of mutual gene pairs in PSN and GRN individually or together with respect to mutual pair correlations in normal and cancer tissues (See Methods). Using MGPNet we could analyze the mutual neighborhoods of PP, PN or NN pairs by selecting genes according to their ranks in terms of connectivity in the descending order and visualizing the pairs’ direct neighborhood of PP, NN or PN type interactions.

We focused on the top ranking negatively correlated NN pairs in normal tissues at protein level for PSN (Supplementary Tables 2-3). As it can be seen in MGPNet, SNCA, which has been shown to be interacting with proteasome (Synder et al, 2003), was a hub and demonstrated a significant example for the distinct coexpression characteristics of NN pairs, as most of its NN neighbors had positive cancer vs normal protein level correlations; this suggested that negative correlations in normal tissues were lost in cancer (Figure 6A). When the same neighborhood was analyzed at the RNA level, negative coexpression trend was observed for cancer vs normal differential coexpression in contrast to protein level (Figure 6B).

**Figure 6.**
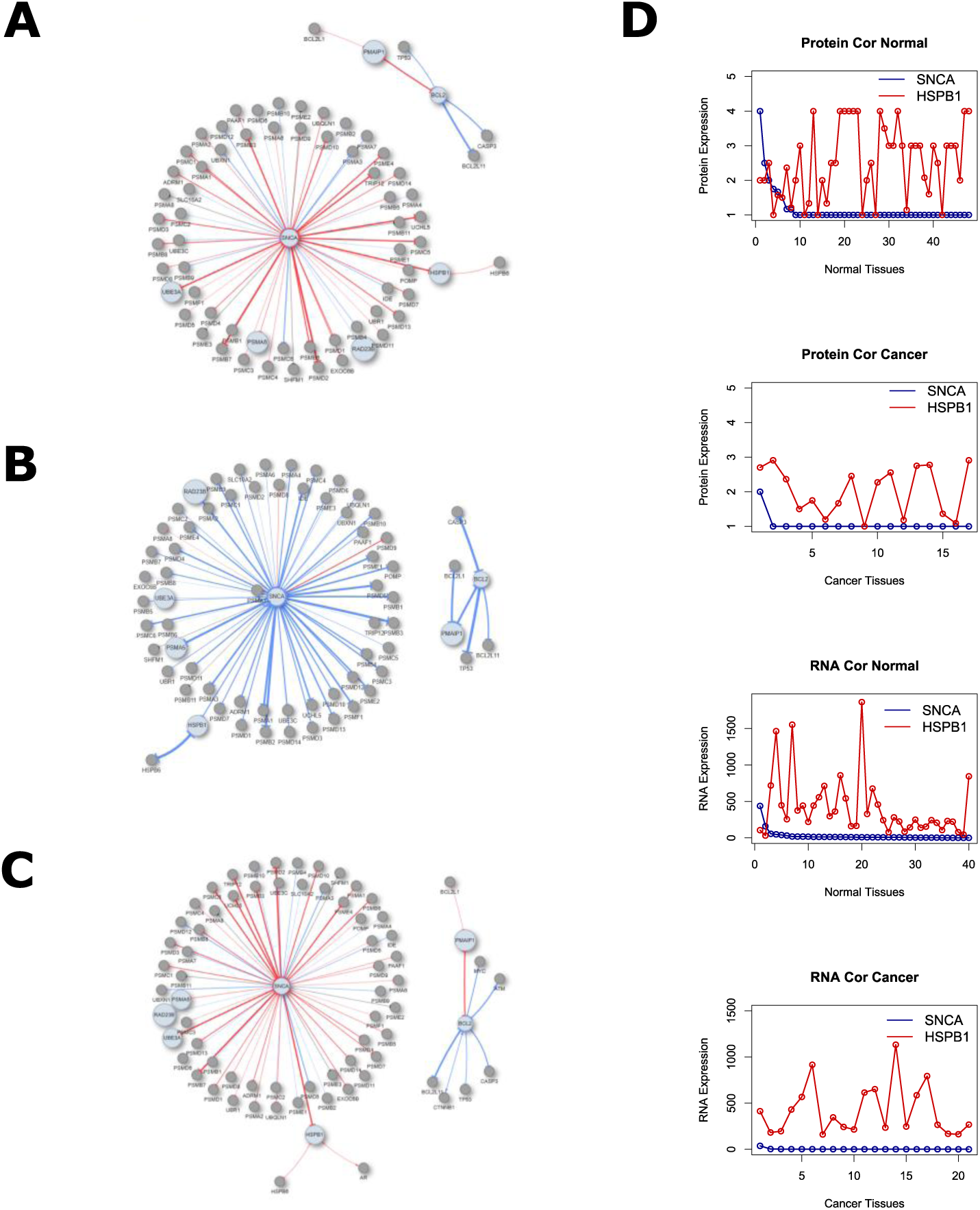
Networks of top ranking negatively correlated NN pairs in PSN visualized by the online tool. A. NN type neighborhood of top ranking negatively correlated NN pairs in PSN visualized by integrating protein coexpression values in cancer vs normal. B. NN type neighborhood of top ranking negatively correlated NN pairs in PSN visualized by integrating RNA coexpression values in cancer vs normal. C. All mutual interactions neighborhood of top ranking negatively correlated NN pairs in PSN visualized by integrating protein coexpression values in cancer vs normal. D. RNA and protein expression values of SNCA in normal or cancer in comparison with HSPB1 values.

We then added the PP and PN type interactions of these top ranking genes so that the entire mutual interaction neighborhood could be observed with integration of the protein coexpression (Figure 6C). There was not a high number of additional interactions and neighbors, thus most of their interactions corresponded to the NN type. Next, we analyzed the protein and RNA expression values of SNCA and its NN neighbors. We observed that SNCA was not expressed significantly at the protein level, whereas its neighbors were expressed in most normal tissues. However, for a few types of tissues (bone marrow, kidney, soft tissue), where SNCA was expressed at not detected-high range, the expression of its neighbor, HSPB1, was lower than other tissues (Figure 6D). On the other hand, HSPB1 levels were high for several tissues such as bronchus, epididymis, and esophagus where SNCA was always not detected. This negative coexpression tendency could not be observed in cancer since SNCA was not significantly expressed at the protein level in most cancer types. This negative coexpression between SNCA and HSPB1 which was detected at protein level, was not observed at the RNA level (Figure 6D). The direct NN neighbors of SNCA were enriched for ubiquitin-proteasome pathway (enrichment p-value < 8.3e-98). Overall, mutually negative interactions of SNCA with the ubiquitin-proteasome genes were negatively correlated across normal tissues, but not cancer tissues, and this observation could be detected at protein level but not at RNA level.

## Discussion

In this study, we observed that mutual gene pairs have distinct characteristics in large scale cellular molecular networks of different types. NN (double negative feedback) pairs, had significantly different topological and coexpression characteristics than the other mutual or non-mutual pairs. This might be relevant to their potential functional roles in cellular information transfer. Whilst PP configuration could stimulate signal amplification and robustness, NN motif could play a role in switch like behavior, as it was reported in a transcription factor-miRNA network and a synthetic gene regulatory network (Cai et al. 2013, Gardner et al. 2000). Switch like behavior supported by NN motif was also shown to play a role in cancer (Yang et al. 2013, Sankpal et al. 2017). Thus, our observation for the significant enrichment of NN, but not PP pattern in human molecular networks, could suggest an enrichment for switch like behavior potential.

Previous statistical investigation of mutual motif structures was limited and did not include two different human molecular networks (Mazurie et al. 2005, Wang et al. 2014, Borotkanics and Lehmann 2015). We were able to confirm some of our observations in two unrelated comprehensive networks. For instance, a higher number of NN pairs was observed in both networks therefore there could be a cellular preference for using NN motif to regulate important molecular events.

All mutual pair genes were more connected than remaining genes, which could support the critical role for mutual pair networks enriched with such structures. For instance, a switch like decision making could be transmitted to other parts of the network. NN pair genes had less of in-degree type connections, whereas more of out-degree connections than PP or PN pair genes in PSN. On the other hand, NN pairs were more connected than PP or PN in GRN for all types of interactions. Therefore, out type but not in type interactions of NN pair genes were higher than other genes for both PSN and GRN.

Different molecular circuits could work together to have a robust control on cellular events (Rata et al. 2018). Similarly different mutual pairs could also operate cooperatively to achieve important transitions such as tumorigenesis. Unlike PP pairs, NN or PN pair genes showed significant overlap among themselves and with each other in both PSN and GRN, therefore there could be an enrichment for cooperative mechanisms among and/or between NN and PN type motifs in human molecular cellular networks. This also provides evidence for a positive selection for double negative feedback motif (NN) type unlike double positive feedback (PP) type. Tied PP patterns could induce cellular signal amplification and bistability, and tied NN motifs could be important for homeostasis (Kim et al. 2008). Therefore, our analysis also suggested a bias for homeostasis rather than signal amplification.

In addition to structural characteristics, we also aimed to evaluate the importance of mutual patterns for human disease, specifically for cancer. We used not only RNA but also protein expression values of different normal tissues and cancer types from HPA. PP and PN pairs generally had significantly higher RNA and protein correlation values for normal and cancer in both PSN and GRN. This could indicate that these pairs tend to work together in different conditions. It also shows that the categorical immunohistochemistry scoring based protein expression values could be confirmed by RNA expression values.

On the other hand, NN pairs exclusively had differential average RNA and average protein coexpression patterns in cancer vs normal tissues for PSN. They had lower RNA coexpression whereas higher protein coexpression in cancer tissues vs normal. The two networks, PSN and GRN, that we analyzed in this study are based on human data, and PSN is specifically based on cancer data (Li et al. 2012). This could be the reason for the PSN-specific differential coexpression of NN. Consequently, NN pairs of PSN might play a role specifically for cancer or non-cancer conditions.

Negative protein coexpression across normal but not cancer tissues might indicate that two genes of a NN, a double negative feedback loop, might have contrasting expression across some normal samples which could be lacking in cancer. Therefore, the significance of the negative effects of NN pair genes to each other for the important biological processes related to cancer should be further investigated.

In this study we were able to recover one such process as ubiquitin-proteasome pathway genes were enriched in negatively correlated NN genes due to their interaction with SNCA, which mostly had negative protein coexpression dominantly in normal tissues only. Previously an antagonistic effect of SNCA was reported for Parkinson’s disease (Zondler et al. 2017).

Furthermore, higher expression of SNCA was associated with better prognosis of some cancer types (Zhang et al. 2022, Zhou et al. 2022). Proteosome genes were also suggested to be prognostic markers for cancer (Larsson et al. 2022). Therefore, MGPNet allowed us to pinpoint SNCA and its NN pairs, enriched in ubiquitin-proteasome pathway, and the switch of direction in their expression correlation in cancer, which warrants further study. Other use cases can be found in the MGPNet tutorial that range from demonstration of SREBF1-SREBF2-PPARG network for visualizing the differences between a) normal and tumor states as well as bb) RNA and protein levels (Demo 6) and obtaining and pruning of a large immediate early gene set (Bahrami et al. 2016) network in GRN to reveal differences at the RNA and protein level differences between selected nodes and neighborhoods (Demo 7). These use cases successfully demonstrate the ability of MGPNet to help establish novel regulatory activity between genes and their proteins in normal and cancer tissues.

There could be several reasons for the absence of correlation for all mutual pairs; for instance, we limited our analysis to only average expression values of a tissue which ignores condition/sample specific variation of RNA and protein expression values. Therefore, some gene pairs which might have coexpression in a specific condition are missed. Furthermore, since correlation is condition-specific, other correlated pairs might be identified by using different expression datasets. For instance, some other mutual pairs might only be correlated in a time-dependent manner, that can be identified using a time-series expression dataset. In addition, not all mutual pairs could show coexpression because of the threshold effect; the expression of gene has to reach a certain threshold in order to show its positive of negative regulatory effect (Little, 2005). Thus, the expression of some genes might not be in the necessary range for having its positive or negative effect and correlation with its target cannot be observed. Furthermore, some of the activation or inhibition relationships in PSN might not have any influence on RNA and/or protein expression levels. For example, posttranslational modifications such as phosphorylation might not affect RNA or protein expression levels.

## Methods

### Datasets

Two comprehensive human molecular interaction networks were used in this study; the gene regulatory network (GRN) and the protein signaling network (PSN) (Müller-Dott S et al. 2023, Wang 2014). GRN, the CollecTRI-derived regulons, was constructed by the integration of various sources such as public transcriptional interaction databases, literature, and computationally predicted regulatory relationships, hence it has a high coverage of transcription factors and a high predictive power for transcriptional factor activities (Müller-Dott S et al. 2023). A large scale human protein signaling network has been built and manually curated, including activating, inhibitory and physical (undirected) interactions in 2014 (Wang 2014). It integrated curated human cancer signaling networks with the BIOGRID protein-protein interactions (Awan et al. 2007; Cui et al. 2007; Li et al., 2012, Chatr-Aryamontri et al., 2015, Stark et al. 2006, Wang et al. 2014) that have positive, negative and physical interactions with no sign. Only positive (activating) and negative (inhibitory) interactions were used in the current study. Therefore, PSN represented a signed and directed cancer protein signaling network. We selected GRN and PSN for the present study, because they were large-scale, comprehensive and different from each other. Respectively, GRN and PSN had 62393 and 41358 signed directed gene pairs with less than 10% overlap and when combined their coverage was larger.

Normal and tumor RNA and protein expression values were obtained from Human Protein Atlas (HPA) database version 23 (proteinatlas.org, Uhlén et al. 2015, Uhlén et al. 2017, Karlsson et al. 2021). RNA HPA tissue gene data and RNA TCGA cancer sample data were used for normal and cancer tissue RNA expression levels. Unlike the normal tissue dataset, RNA TCGA cancer sample data provided values at the sample level, therefore we first calculated median expression values for multiple samples of a cancer type for each gene. For both normal and cancer RNA datasets, mean values were calculated for multiple TPM/FPKM expression values (genes with more than one Ensembl Gene ID per a Gene symbol) per gene in the same tissue type.

Normal tissue data and pathology data of HPA were used for normal and cancer tissue protein expression levels. For protein expression, immunohistochemistry scoring based categorical data was converted to numbers as follows; ‘Not detected’ is 1, ‘Low’ is 2, ‘Medium’ is 3, and ‘High’ is 4. For normal tissue types with more than one cell type we calculated the mean expression. For cancer types multiple scoring values were provided by HPA unlike normal tissues. Therefore, we calculated mean values for expressions of genes for a particular cancer type, if there were at least 10 different scoring values. For both the normal and cancer protein datasets, mean values were used for genes with more than one Ensembl Gene ID per a Gene symbol.

### Network Analysis

The majority of the interacting gene pairs in both gene regulatory and protein signaling networks do not act mutually. For the ones that do, there are 3 possible structural patterns in signed directed networks; mutually positive (PP), mutually negative (NN) and positive-negative (PN) pairs (Figure 1A). We analyzed the number of occurrences of PP, NN and PN pairs in both PSN and GRN. For significance testing, we randomized the networks by shuffling the positive and negative signs of all interacting pairs. Therefore, the gene pairs and the number of positive and negative signs were kept same with the original network but the signs of the gene pairs were randomized.

For connectivity, total degree values which are the total number of interactions (or neighbors) of a gene were analyzed. In addition, different types of degree values with respect to the type of interaction were also evaluated; in, out, positive, negative, in-positive, in-negative, out positive and out negative degree, defined based on the direction (in or out) and sign (positive or negative) of the interactions. For example, if a gene only participates in a single NN type pattern and does not have any other interaction, it will have an in degree of 1, out degree of 1, positive degree of 0, negative degree of 2, negative in degree of 1, negative out degree of 1, and a total degree of 2. The significance of the difference between the two different degree value sets was calculated by a Wilcoxon signed-rank test.

We also analyzed the intersection (co-occurence) of different mutual pairs; i.e, calculated as the number of cases where a particular mutual subnetwork shares a common gene with a particular other mutual subnetwork (PP, PN, or NN). For instance, if a gene is involved in one NN and two PP type motifs, there are two NN-PP intersection values. All combinations of mutual pair intersections were analyzed; PP-PP, PN-PN, NN-NN, PP-PN, PP-NN, and PN-NN. We calculated total co-occurences for all genes for the real and randomized networks. For 1000 random iterations co-ccurences were calculated and the number of times their values were higher/lower or equal to the real network value was used a randomization p-value. Co-occurence values with a p-value less than 0.05 were considered significant.

### RNA and Protein Coexpression Analysis

HPA RNA and protein expression datasets were integrated to mutual gene pairs from GRN and PSN via Official Gene Symbols. For two genes, the same set of tissues, or cancer types were selected with their expression values to calculate the Spearman’s correlations. Spearman correlation p-values and False Discovery Rate (FDR) values for a particular list of genes (i.e,, PP gene pairs) were also calculated. Such correlation parameters were calculated for both networks including their PP, NN and PN pairs, as well as randomized networks and were compared for testing significance. For 1000 iterations, median Spearman correlation rho values of random sets of pairs with the same signed interactions (only positive interactions for PP, only negative interactions for NN, positive and negative interactions for PN) were calculated and the fraction of those equal to or higher/lower than the real set’s median rho value was used as a randomization based p-value. In addition, the significance of differences between two sets of coexpression values was tested by Kolmogorov–Smirnov and Wilcoxon signed-rank tests. P-values less than 0.05 were considered significant. R scripts for the analysis of network level characteristics and coexpression analysis are available at https://github.com/ertuda.

### Mutual Pairs Network Analysis Online Tool

We constructed an online tool, MGPNet, Mutual Gene Pairs Network, to allow visualization of the mutual interactions of a gene or genes of interest by the user. Nonmutual interactions were omitted from MGPNet because of the large-scale of the data hence the difficulty of visualization. MGPNet is a user-friendly Shiny web application to deploy our data of gene-gene interaction pairs from GRN and PSN both in table and network formats (Chang et al. 2024). It enables users to generate networks via *visNetwork* package in R (Almende et al. 2022) so that the direction of interaction can be visualized by an arrow (positive) or a bar (negative) over an edge in GRN, PSN or a combination of GRN and PSN. The networks could be pruned based on user defined selections of PP, PN, NN interactions for genes/proteins and the incremental thresholds, i.e., 0.1, put on the edges based on the values of coexpression from HPA. Moreover, normal or cancer tissue correlation values from HPA as well as their paired differences could be used to color the edges (red positive; blue negative correlations) while the edge weight refers to the strength of the correlation. Interaction data of mutual gene pairs reflects both mutuality types (PP, PN or NN) and correlation values with p- and FDR-values in normal and cancer status at the RNA and protein levels. MGPNet provides the gene names for user selection in the order of their ranks according to the number of connections (degrees) they have. This enables the user to select the direct neighborhood of most highly connected genes by clicking in the given order. MGPnet has multiple filters, e.g., thresholds for expression correlation and number of degrees. User specific filtering results in in customized and publication quality networks, which can be annotated by gene set enrichment based on Reactome terms (Yu et. al., 2016). MGPNet can be accessed at konulabapps.bilkent.edu.tr:3838/MGPNet/, on which a detailed tutorial and additional use cases (Demo 1 to Demo 7) can be found. R scripts for the analysis of network level characteristics and coexpression analysis are available at.

## Author Contributions

ED and OK designed the study, ED performed the statistical network and correlation analyses and wrote the scripts for those purposes; OK and ED conceptually developed the online application; MVO implemented it in R Shiny server; OK and MC further developed the application, wrote the tutorial and relevant parts of the article. ED wrote and OK revised and edited the manuscript.

## Acknowledgments

ED is Chief Executive Officer of Bomemod Y.M.S.T. Ltd. Sti. His role in the company did not interfere with the design and implementation of the study. This study has partly been funded by the European Horizon’s research and innovation program HORIZON-HLTH-2022-STAYHLTH-02 under agreement No 101095679 (to OK). Funded by the European Union. Views and opinions expressed are however those of the author(s) only and do not necessarily reflect those of the European Union. Neither the European Union nor the granting authority can be held responsible for them.

## Supplementary Text

### Correlation Analysis of PP and PN pair genes

PP pairs had significantly higher RNA correlation values than other positively interacting pairs across normal tissues and cancer types in both PSN and GRN (Supplementary Figure 1, PSN: Kolmogorov–Smirnov test p value 1.549e-11 for normal and 0.0003277 for cancer, Wilcox test p value 1.091e-13 for normal and 1.602e-05 for cancer, GRN: Kolmogorov–Smirnov test p value 5.107e-14 for normal and 0.000351 for cancer, Wilcox test p value < 2.2e-16 for normal and 1.569e-05 for cancer) (Supplementary Figure 1). They also had significantly higher correlation values than random positively interacting pairs of the same size (random sampling test p value 0.001 for normal and 0.002 for cancer in PSN, 0.001 for normal and 0.001 for cancer in GRN).

Similar to PP pairs, PN pairs also had significantly higher RNA correlation values than other positively or negatively interacting pairs in both networks (Supplementary Figure 2, PSN: Kolmogorov–Smirnov test p value 0.0272 for normal and 0.02748 for cancer, Wilcox test p value 0.003932 for normal and 0.01815 for cancer, GRN: Kolmogorov–Smirnov test p value 2.289e-05 for normal and 0.02132 for cancer, Wilcox test p value 4.375e-05 for normal and 0.005681 for cancer) (Supplementary Figure 1). They also had significantly higher RNA correlation values than random positively interacting pairs of the same size (random sampling test p value 0.008 for normal and 0.05 for cancer in PSN, 0.001 for normal and 0.04 for cancer in GRN) (Supplementary Figure 1).

PP pairs had significantly higher protein correlation values across normal tissues than other positively interacting pairs in PSN, but not across cancer types (Supplementary Figure 1, PSN: Kolmogorov–Smirnov test p value 0.001955 for normal and 0.351 for cancer, Wilcox test p value 5.034e-05 for normal and 0.1983 for cancer). PP pairs had significantly higher protein correlation values for both normal and cancer in GRN (Kolmogorov–Smirnov test p value 0.0006726 for normal and 2.197e-06 for cancer, Wilcox test p value 0.002114 for normal and 2.601e-08 for cancer) (Supplementary Figure 1). They also had significantly higher correlation values than random positively interacting pairs of the same size (random sampling test p value 0.026 for normal and 0.17 for cancer in PSN, 0.007 for normal and 0.001 for cancer in GRN).

PN pairs had significantly higher protein expression across cancer types but normal tissues in both networks (Supplementary Figure 2, PSN: Kolmogorov–Smirnov test p value 0.2262 for normal and 0.005976 for cancer, Wilcox test p value 0.1439 for normal and 0.01132 for cancer, GRN: Kolmogorov–Smirnov test p value 0.1327 for normal and 0.04636 for cancer, Wilcox test p value 0.03683 for normal and 0.02314 for cancer) (Supplementary Figure). They also had significantly higher correlation values than random positively interacting pairs of the same size (random sampling test p value 0.037 for normal and 0.002 for cancer in PSN, 0.031 for normal and 0.024 for cancer in GRN).

Frequently differentially more correlated genes (with at least 4 different interactions with a positive RNA or protein differential correlation in cancer vs normal) of PP and PN pairs were analyzed furthermore. The PN neighborhood of frequently differentially more correlated PN pair genes; CDK1 and CCNB1 in PSN, and E2F1 in GRN, were displayed in MGPNet (Supplementary Figure 2A-B). The PP neighborhood of frequently differentially more correlated PP pair genes; BLK, EPHB2, and FGR in PSN, and TFAP2A, MYC, SP1, and TP53 in GRN, were displayed in MGPNet (Supplementary Figure 2C-D).

---

**Supplementary Figure 1.**
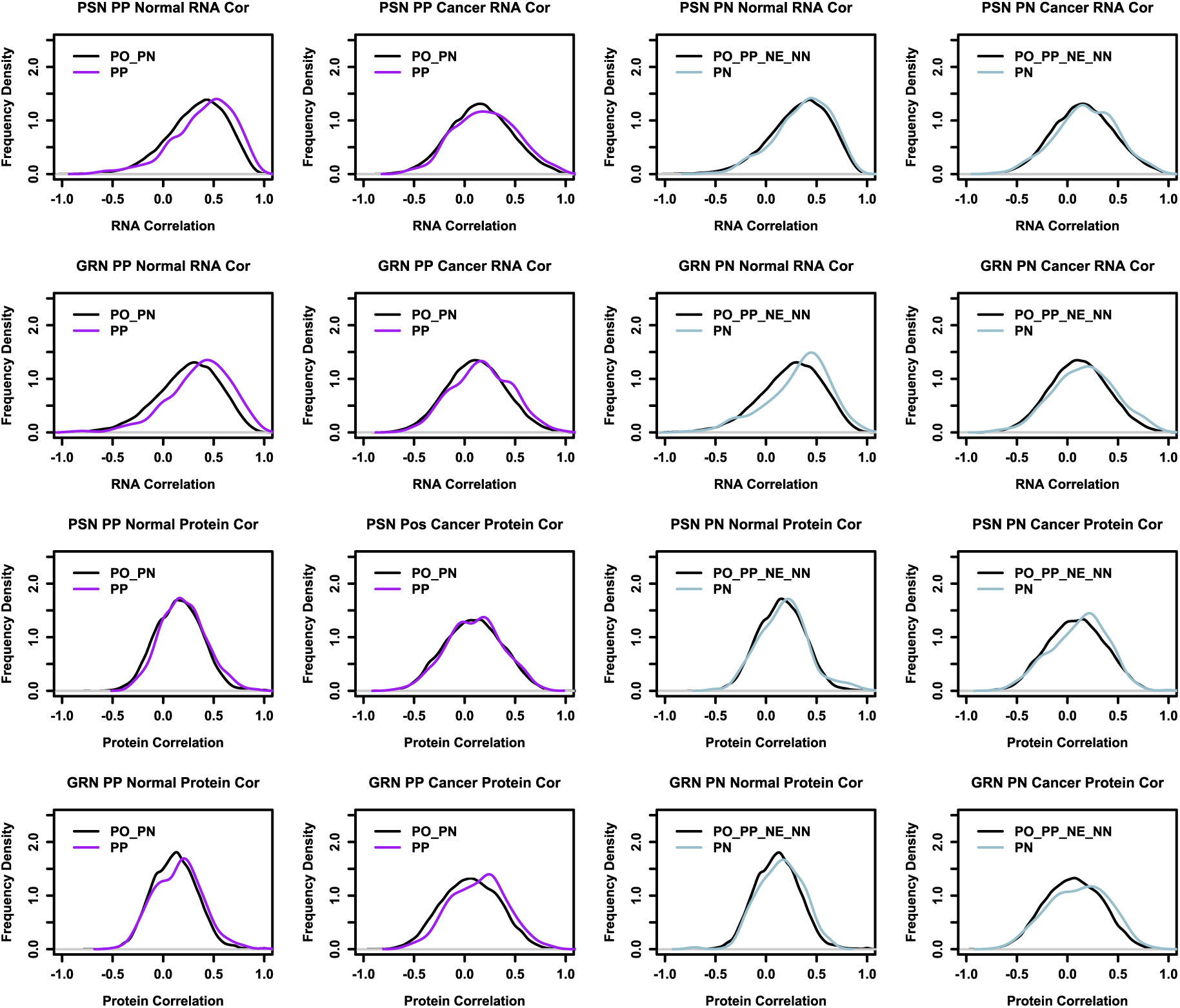
RNA and protein expression correlation distribution of PP and PN pair genes with comparison to other genes that participate in positive or negative interactions in PSN and GRN. Kernel density plots of PP and PN pairs’ correlation values together with the remaining genes with positive/negative interactions; non-mutual negative interaction pairs (PO), non-mutual negative interaction pairs (NE), PP, NN and PN pairs. Correlation (cor) was measured by Spearman’s rho. Correlation analysis included normal and cancer specific expression values.

**Supplementary Figure 2.**
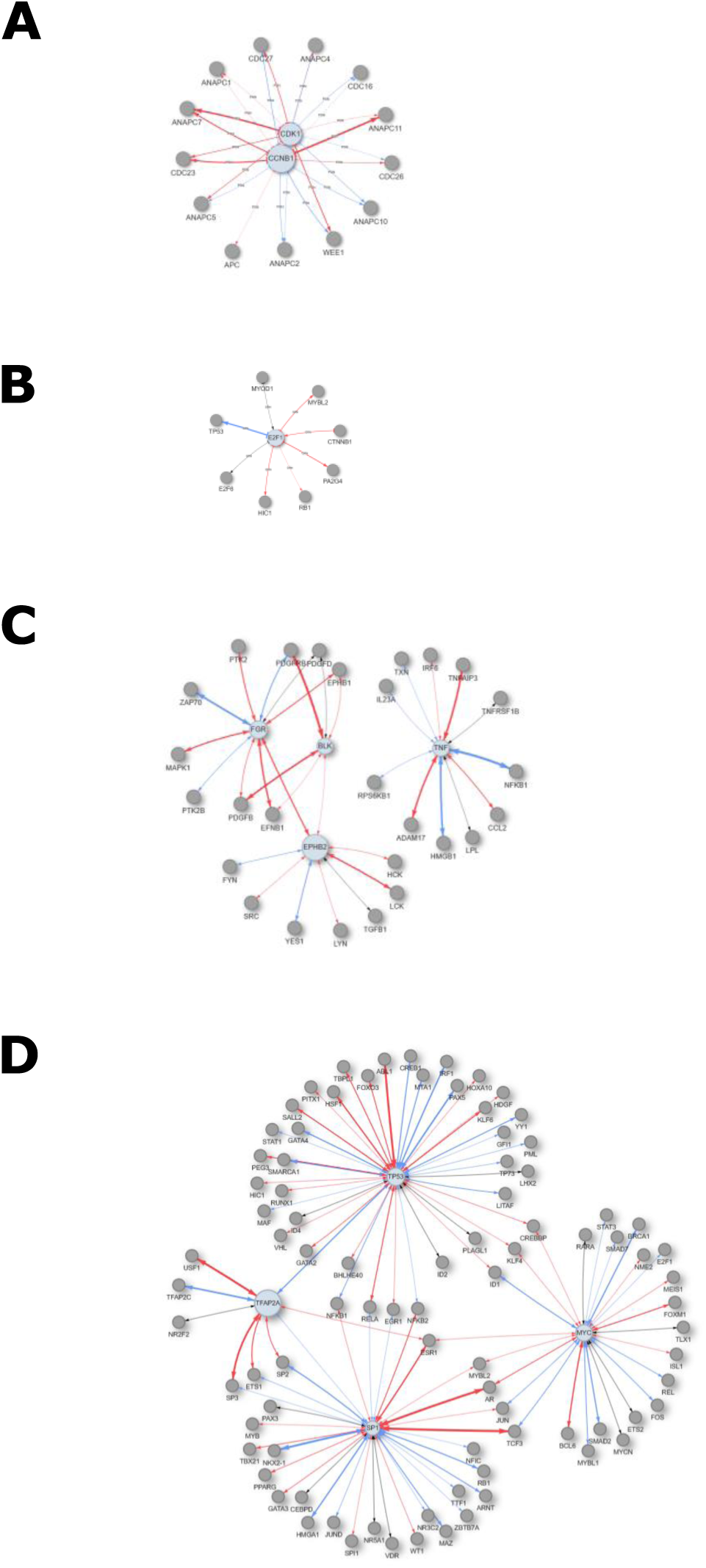
Mutual neighborhood of frequently highly correlated PP and PN pair genes for A. PN pair genes in PSN, B. PN pair genes in GRN, C. PP pair genes in PSN, and D. PP pair genes in GRN.

